# Peripheral Blood Single-Cell Sequencing Uncovers Common and Specific Immune Aberrations in Fibrotic Lung Diseases

**DOI:** 10.1101/2023.09.20.558301

**Authors:** Amy Y. Zhao, Avraham Unterman, Nebal Abu Hussein, Prapti Sharma, Jasper Flint, Xiting Yan, Taylor S. Adams, Aurelien Justet, Tomokazu S. Sumida, Jiayi Zhao, Jonas C. Schupp, Micha Sam B. Raredon, Farida Ahangari, Yingze Zhang, Ivette Buendia-Roldan, Ayodeji Adegunsoye, Anne I. Sperling, Antje Prasse, Changwan Ryu, Erica Herzog, Moises Selman, Annie Pardo, Naftali Kaminski

## Abstract

**Rationale and Objectives:** The extent and commonality of peripheral blood immune aberrations in fibrotic interstitial lung diseases are not well characterized. In this study, we aimed to identify common and distinct immune aberrations in patients with idiopathic pulmonary fibrosis (IPF) and fibrotic hypersensitivity pneumonitis (FHP) using cutting-edge single-cell profiling technologies.

**Methods:** Single-cell RNA sequencing was performed on patients and healthy controls’ peripheral blood and bronchoalveolar lavage samples using 10X Genomics 5’ gene expression and V(D)J profiling. Cell type composition, transcriptional profiles, cellular trajectories and signaling, and T and B cell receptor repertoires were studied. The standard Seurat R pipeline was followed for cell type composition and differential gene expression analyses. Transcription factor activity was imputed using the DoRothEA-VIPER algorithm. Pseudotime analyses were conducted using Monocle3, while RNA velocity analyses were performed with Velocyto, scVelo, and CellRank. Cell-cell connectomics were assessed using the Connectome R package. V(D)J analyses were conducted using CellRanger and Immcantation frameworks. Across all analyses, disease group differences were assessed using the Wilcoxon rank-sum test.

**Measurements and Main Results:** 327,990 cells from 83 samples were profiled. Overall, changes in monocytes were common to IPF and FHP, whereas lymphocytes exhibited disease-specific aberrations. Both diseases displayed enrichment of CCL3^hi^/CCL4^hi^ CD14+ monocytes (p<2.2e-16) and S100A^hi^ CD14+ monocytes (p<2.2e-16) versus controls. Trajectory and RNA velocity analysis suggested that pro-fibrotic macrophages observed in BAL originated from peripheral blood monocytes. Lymphocytes exhibited disease-specific aberrations, with CD8+ GZMK^hi^ T cells and activated B cells primarily enriched in FHP patients. V(D)J analyses revealed unique T and B cell receptor complementarity-determining region 3 (CDR3) amino acid compositions (p<0.05) in FHP and significant IgA enrichment in IPF (p<5.2e-7).

**Conclusions:** We identified common and disease-specific immune mechanisms in IPF and FHP; S100A^hi^ monocytes and SPP1^hi^ macrophages are common to IPF and FHP, whereas GMZK^hi^ T lymphocytes and T and B cell receptor repertoires were unique in FHP. Our findings open novel strategies for the diagnosis and treatment of IPF and FHP.

## Introduction

While idiopathic pulmonary fibrosis (IPF) and fibrotic hypersensitivity pneumonitis (FHP) are often hard to distinguish, their etiologies and immune mechanisms are presumed to be drastically different [1]. IPF, the prototypical fibrotic lung disease exhibiting a usual interstitial pneumonia (UIP) pattern by histology and imaging, largely does not have an apparent causative injury, is attributed to epithelial activation, fibroblast accumulation, and aberrant remodeling in the context of aging, telomere attrition and genetic predisposition [2]. On the other hand, FHP is attributed to a non-resolving inflammatory response to exposure to an inhaled foreign antigen [3]. As mentioned, the diseases are often indistinguishable and even confused clinically [4]. They share similar genetic architecture, association with telomere attrition and tissue gene expression patterns [5–9].

It has long been recognized that IPF and FHP may exhibit distinct immune profiles. Indeed, previous studies have demonstrated that transcriptional differences in peripheral blood mononuclear cells (PBMCs) could indicate disease presence, prognosis, and potential mechanisms of fibrotic lung diseases [10–16]. Peripheral monocyte count correlates with worse patient outcomes in IPF and other interstitial lung diseases [12, 17, 18]. The IPF Cell Atlas, a resource that has vastly expanded single-cell data sharing, and other studies have shown that SPP1^hi^ macrophages are significantly expanded in fibrotic lung disease [19–21]. Lymphoid cells are also dysregulated in IPF and FHP. In IPF, decreased *CXCR3* and increased *CCR4* expression on bronchoalveolar lavage (BAL) T cells suggest a Th1 to Th2 shift [22]. FHP patients exhibit higher CD4+ to CD8+ T cell ratios, decreased gamma-delta T cells (Tγδ), and increased memory CD4+ to CD8+ T cells. However, functional impairments are reported in the lymphocytes of FHP patients, with CD8+ T cells losing their effector function and CD4+ cells showing a Th2 predominance with increased *CXCR4* expression [23]. While the above studies highlighted the importance of peripheral immune cells in both diseases, as well as potential differences, a deep, single cell resolution analysis of common and disease specific immune mechanisms in IPF and FHP is still missing.

To address these gaps, we performed single-cell RNA sequencing on PBMC and BAL samples obtained from individuals with IPF and FHP. We identified specific immune signatures associated with the different disease subtypes. Both lung diseases showed an increase in S100A^hi^ and CCL3/CCL4^hi^ CD14+ monocytes, and trajectory analyses suggested that S100A^hi^ monocytes may give rise to SPP1^hi^ alveolar macrophages. Additionally, we found a distinct subpopulation of CD8+ T cells that was more abundant in patients with FHP. Abnormal cell-to-cell signaling was also observed in both diseases compared to healthy controls. Lymphocyte receptor repertoire analyses revealed distinct compositions in IPF and FHP. Finally, an online data sharing, mining and dissemination portal based on the IPF Cell Atlas accompanies this article.

## Methods

### Study Patients and Samples

Peripheral blood mononuclear cells (PBMCs) and bronchoalveolar lavage (BAL) samples were obtained from patients and healthy controls. PBMC samples were collected from 46 patients with idiopathic pulmonary fibrosis (IPF), 14 patients with fibrotic hypersensitivity pneumonitis (FHP), 5 patients with non-fibrotic hypersensitivity pneumonitis (non-FHP), and 31 healthy controls. BAL samples were obtained from 10 patients with IPF, 6 FHP patients, and 4 non-FHP patients. The diagnosis of FHP and non-FHP was based on the documentation of bird exposure by clinical history and serum-specific immunoglobulins, by the presence of typical imaging findings on high-resolution computed tomography, and BAL lymphocytosis. In two FHP patients, diagnosis was confirmed by histopathology findings. The diagnosis of IPF was established based on the presence of typical/probable UIP either by HRCT and/or lung morphology in the appropriate clinical setting. Before processing the samples, a multidisciplinary diagnostic team corroborated all final diagnoses based in the more recent international guidelines [6, 7]. Biospecimens and corresponding clinical data were obtained from the Yale Lung Repository (TYLR, IRB protocol ID 1307012431) housed within the ILD Center of Excellence at the Yale School of Medicine, and from Mexico (IRB protocol 2000030096; National Institute of Respiratory Diseases (INER) G54-15). Biospecimens from healthy subjects without known inflammatory or lung disease were also obtained from TYLR and G54-15.

### Isolation and Cryopreservation of PBMCs

PBMCs were isolated from whole blood using density gradient centrifugation, washed, resuspended, and cryopreserved in 10% DMSO in heat inactivated FBS for long-term storage.

### PBMC and BAL Sample Preparation, Sequencing, and Pre-Processing

This section’s methods were adapted from Unterman *et al*. [16]. Cryopreserved samples were thawed, and PBMCs/BAL cells were processed following the manufacturer’s protocols (10X Genomics, CG00039). Cell concentration was determined, and single-cell libraries were constructed using 10X Chromium technology (10X Genomics, CG000331). The cDNA libraries were sequenced by an Illumina HiSeq4000 (150 million reads per sample). Sequencing reads were demultiplexed and aligned to the GRCh38 reference genome (CellRanger 2020-A) using CellRanger v7.0.0. Quality control steps were performed to filter out low-quality cells. Data were normalized, scaled, and clustered using *Seurat*. Two patient samples were identified as having monoclonal B cells and thus removed from downstream analyses. Moreover, any doublets were removed. Differential composition and gene expression analyses were performed to identify disease-specific and shared changes.

### Transcription Factor Activity Imputation

Transcription factor (TF) activity was imputed using the DoRothEA-VIPER algorithm [24]. TF activity scores were calculated based on target gene expression, and heatmaps were generated to visualize TF activity patterns. Differential TF activities between disease subtypes were assessed using the Wilcoxon rank-sum test.

### Trajectory Analysis

Pseudotime analysis was conducted using Monocle3 to infer cellular trajectories and identify temporally differentially expressed genes. RNA velocity analysis was conducted using Velocyto, scVelo, and CellRank to estimate transcriptional dynamics and predict cellular fates.

### Cell-Cell Signaling Analysis

The R package *Connectome* was used to assess cell-cell connectomics within each disease condition using a curated version of the FANTOM5 database of ligand-receptor interactions [25, 26].

### B and T Cell Receptor Repertoire Analysis

T and B cell receptor libraries were processed using the CellRanger v7.0.0 V(D)J pipeline to assemble contigs and annotate V(D)J segments. The Immcantation Framework was used for re-annotation and downstream analysis. Clonal groups were defined based on shared gene annotations and junction lengths. Clonal abundance and diversity were assessed using the Alakazam package. Expanded clonal lineages were identified based on fractional abundance and the presence of at least five corresponding cells. Differences between disease groups were evaluated using the Wilcoxon rank-sum test. Amino acid frequencies in the CDR-H3 segment were calculated to determine antigen-binding region chemistries.

### Data Availability

Feature-barcode matrices and Seurat object will be deposited in GEO. An accompanying web server, ILD Immune Cell Atlas, is created and maintained by Jasper Flint and Prapti Sharma and will be available for data browsing (http://ildimmunecellatlas.com).

Additional details can be found in the online data supplement.

## Results

### Single-Cell RNA-Sequencing Reveals Distinct, Known Peripheral Blood Cell Populations

96 PBMC samples (31 healthy controls, 14 FHP, 5 non-FHP, and 46 IPF) were collected at or near time of diagnosis for single-cell RNA sequencing processing by 10X Genomics 5’ kits (**Figure 1A; Table E1**). Samples were matched closely in age, sex, ethnicity, smoking history, and pulmonary function tests. Ethnicity did not significantly differ across disease cohorts (p>0.19). Age and sex showed no significant differences between IPF, FHP, and controls (p>0.18). Non-FHP had younger females compared to other disease and control groups (p<0.05). There were no significant differences in smoking status across groups (p>0.10). Percent predicted forced vital capacity (FVC) and diffusing capacity of the lungs for carbon monoxide (DLCO) differed significantly between control and fibrotic disease (p<0.05) but not between IPF versus FHP (p>0.30).

**Figure 1.**
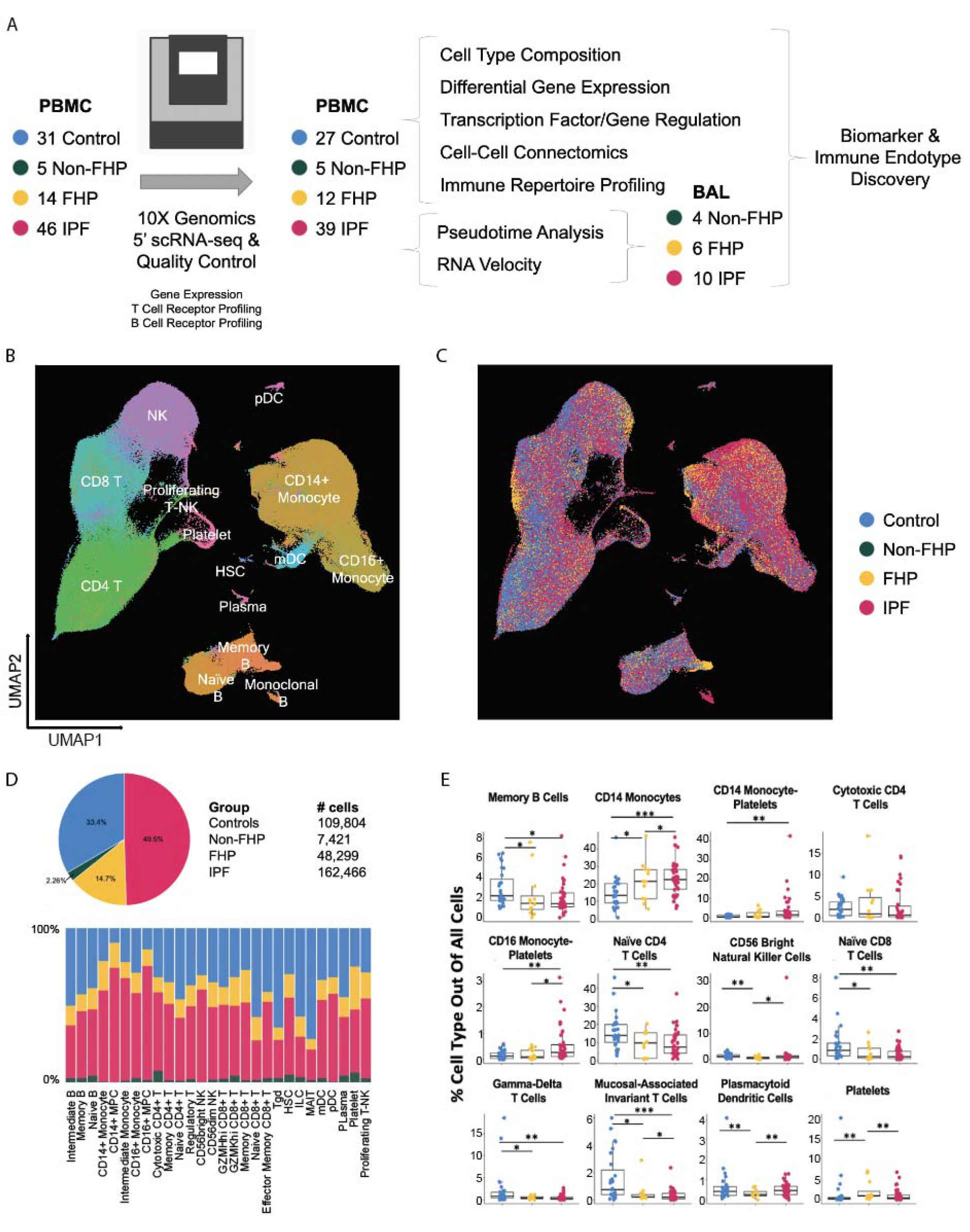
Study Design and PBMC Cell Type Proportions. **(A)** Cryopreserved PBMCs from 46 IPF patients, 14 FHP patients, 5 non-FHP patients, and 31 healthy controls underwent 10X Genomics 5’ single-cell RNA sequencing in randomized processing batches. After filtering, 83 samples (27 controls, 12 fibrotic HP, 5 non-fibrotic HP, and 39 IPF) remained for downstream analysis of cell type composition, gene expression, cell-cell communication, and T and B cell receptor repertoires. Additionally, BAL samples from 10 IPF patients, 6 FHP patients, and 4 non-FHP patients were processed. BAL myeloid cells were combined with those of PBMCs for pseudotime and RNA velocity analyses. UMAP representation of 327,990 PBMCs after quality control, labeled by **(B)** all major cell types and **(C)** disease subtypes. **(D)** Cell type composition by disease or healthy control group. **(E)** Percentage of cell types for each sample, grouped by disease subtype. *p<=0.05, **p<=0.01, ***p<=0.001. PBMC: peripheral blood mononuclear cell; BAL: bronchoalveolar lavage; UMAP: uniform manifold approximation and projection; IPF: idiopathic pulmonary fibrosis; FHP: fibrotic hypersensitivity pneumonitis.

After rigorous quality control and filtering, 327,990 cells from 83 samples (27 controls, 12 FHP, 5 non-FHP, and 39 IPF) remained **(Table 1)**. The cell distribution included 109,804 cells from healthy controls, 48,299 from FHP, 7,421 from non-FHP, and 162,466 from IPF. Manual annotation based on gene expression profiles and cross-referencing with *Azimuth* and *SingleR* allowed the identification of major peripheral mononuclear blood cell types, such as monocytes, CD4+ T cells, CD8+ T cells, Natural Killer (NK) cells, B cells, dendritic cells. Additional cell types, including hematopoietic stem cells (HSCs) and platelets, were also detected in patient samples (**Figure 1A-D)**.

**Table 1.** Summary of demographic and clinical characteristics. Summarized information for patients whose single-cell RNA-sequencing PBMC data were analyzed. Abbreviations: IPF: idiopathic pulmonary fibrosis; FHP: fibrotic hypersensitivity pneumonitis. FVC: Forced Vital Capacity; DLCO: Diffusing Capacity of Carbon Monoxide; SD: Standard Deviation

|  | **CONTROL (27)** | **NON-FHP**<br>**(5)** | **FHP**<br>**(12)** | **IPF**<br>**(39)** |
| --- | --- | --- | --- | --- |
| **AGE**<br>(MEAN ± SD) | 69.30 ± 6.36 | 47.00 ± 13.55 | 67.25 ± 11.05 | 69.49 ± 7.70 |
| **SEX** |  |  |  |  |
| **MALE** | 17 | 0 | 5 | 29 |
| **FEMALE** | 10 | 5 | 7 | 10 |
| **ETHNICITY** |  |  |  |  |
| **CAUCASIAN** | 18 | 0 | 7 | 6 |
| **HISPANIC** | 9 | 5 | 5 | 32 |
| **OTHER** | 0 | 0 | 0 | 1 |
| **SMOKING** |  |  |  |  |
| **NEVER** | 7 | 2 | 7 | 12 |
| **EVER** | 15 | 3 | 5 | 27 |
| **NO DATA** | 5 | 0 | 0 | 0 |
| **% FVC**<br>(MEAN ± SD) | 93.77 ± 16.62 | 75.00 ± 15.63 | 63.22 ± 17.72 | 72.62 ± 11.72 |
| **% DLCO**<br>(MEAN ± SD) | 106.33 ± 12.71 | 61.00 ± 16.17 | 42.33 ± 15.34 | 47.08 ± 13.61 |

### Peripheral Blood Mononuclear Cells Demonstrate Significant Disease-Specific and Shared Fibrosis Cell Type Proportional Differences

Once annotated, the proportions of cell types in each patient sample were calculated (**Figure 1E**). IPF and FHP showed similar trends compared to controls in monocytes, B, and T cell subtypes. Specifically, IPF and FHP had decreased proportions of memory B cells (IPF average: 2.0%; FHP: 2.2%; control: 3.0%; IPF vs control: p=0.011; FHP vs control: p=0.045), naïve CD4 T cells (IPF average: 9.6%; FHP: 9.5%; control: 16%; IPF vs. control: p=0.003; FHP vs. control: p=0.05), naïve CD8 T cells (IPF average: 0.52%; FHP: 0.66%; control: 1.4%; IPF vs. control: p=0.001; FHP vs. control: p=0.04), mucosal-associated invariant T (MAIT) cells (IPF average: 0.31%; FHP: 0.49%; control: 1.6%; IPF vs. control: p=6.2e-5; FHP vs. control: p=0.008), and Tγδ cells (IPF average: 0.50%; FHP: 0.49%; control: 2.2%; IPF vs. control: p=0.005; FHP vs. control: p=0.04). Conversely, CD14+ monocytes were increased in fibrotic conditions versus control (IPF average: 23%; FHP: 22%; control: 15%; IPF vs. control: p=0.00017; FHP vs. control: p=0.048).

Disease-specific shifts in cell composition were also observed. FHP had higher platelet proportions than healthy controls and IPF (IPF average: 1.1%; FHP: 2.1%; control: 1.4%; FHP vs. control: p=0.002; FHP vs. IPF: p=0.02). Additionally, FHP showed decreased proportions of CD56^bright^ NK cells (IPF average: 1.8%; FHP: 0.76%; control: 1.4%; FHP vs. control: p=0.008; FHP vs. IPF: p=0.01) and plasmacytoid DCs (IPF average: 0.50%; FHP: 0.31%; control: 0.63%; FHP vs. control: p=0.003; FHP vs. IPF: p=0.01). In IPF, CD14+ monocyte-platelet (IPF average: 4.0**%;** control: 0.70%; IPF vs. control: p=0.0016) and CD16+ monocyte-platelet complexes (IPF average: 0.53%; FHP: 0.18%; control: 0.17%; IPF vs. control: p=0.004; IPF vs. FHP: p=0.02) were increased. Non-FHP patients had increased cytotoxic CD4+ T cells compared to controls (NFHP average: 7.7%; control: 2.4%; NFHP vs control: p=0.04) (**Figure E1**).

The proportions of CD14+ monocytes, CD16+ monocytes, and intermediate monocytes, were not significantly different among IPF, FHP, and healthy controls **(Figure E2)**. As a proportion of CD4+ T cells, naïve cells were decreased in FHP and IPF compared to controls (IPF average: 41%; FHP: 36%; control: 51%; IPF vs. control: p=0.04; FHP vs. control: p=0.04), while memory cells were increased in IPF compared to controls (IPF average: 39%; control: 32%; IPF vs. control: p=0.03). FHP showed higher proportions of memory cells out of CD4+ T cells than non-FHP (FHP average: 39%; NFHP: 35%; FHP vs. NFHP: p=0.04). Regulatory T cells as a proportion of CD4+ T cells were increased in IPF compared to controls (IPF average: 7.1%; control: 5.7%; IPF vs. control: p=0.03). Among CD8 T cells, memory cells (IPF average: 14%; control: 7.7%; IPF vs. control: p=0.03) were increased, while MAIT cells (IPF average: 3.8%; control: 10%; IPF vs. control: p=0.02) were decreased in IPF compared to controls **(Figure E3)**. Within B cells, memory B cells were lower (FHP average: 20%; control: 30%; IPF vs. control: p=0.01), and naïve B were higher (FHP average: 69%; control: 57%; IPF vs. control: p=0.05) in FHP compared to controls **(Figure E4)**.

### CD14+ Monocytes Harbor Shared Fibrotic Niche

To improve the resolution of the molecular events in monocytes, we subsetted and reclustered 94,967 monocytes and monocyte-platelet complexes **(Figure 2A)**. The different cell types were identified based on highly expressed genes (*CD14, FCGR3A, PF4/PPBP* for CD14+ monocytes, CD16+ monocytes, and monocyte-platelet complexes, respectively) **(Figure E5)**. CD14+ monocytes were visualized by disease subtype, revealing disease-specific shifts with over-representation of cell clusters in FHP and IPF compared to non-FHP and healthy controls **(Figure 2B-C)**.

**Figure 2.**
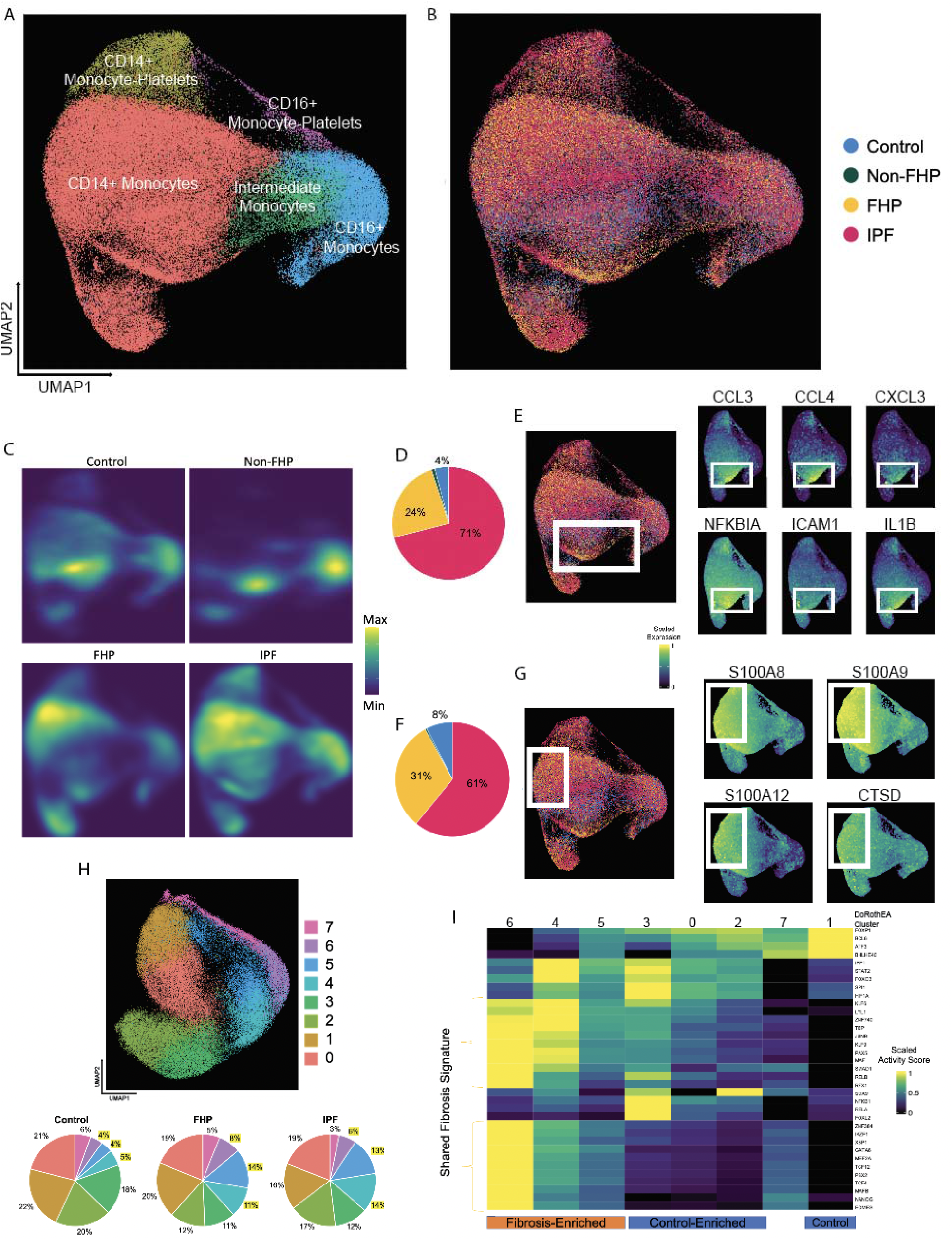
Monocyte Subpopulations Enriched in Fibrosis. UMAP representation of 94,967 cells by **(A)** monocyte subtypes and **(B)** disease subtypes. **(C)** Density graph of monocytes by disease subtype. **(D)** Breakdown of fibrosis-enriched *CCL3/CCL4hi* monocytes (p<2.2e-16) by disease. **(E)** Gene expression of *CCL3, CCL4, CXCL3, NFKBIA, ICAM1,* and *IL1B*. **(F)** Breakdown of fibrosis-enriched *S100Ahi* monocytes (p<2.2e-16) by disease. **(G)** Gene expression of *S100A8, S100A9, S100A12,* and *CTSD.* **(H)** UMAP of monocytes clustered by DoRothEA-VIPER transcription factor activity scores, and the breakdown of the DoRothEA-based clusters. Cluster 5 is significantly expanded in IPF and FHP versus controls (p=0.040; p=0.024). **(I)** Heatmap showing TFs with top variable imputed activities across monocyte DoRothEA clusters, with hierarchically clustered rows and columns. UMAP: uniform manifold approximation and projection; IPF: idiopathic pulmonary fibrosis; FHP: fibrotic hypersensitivity pneumonitis; TF: transcription factor.

Two CD14+ monocyte subtypes – one enriched in CCL3^hi^/CCL4^hi^ and another in S100Ahi-expressing monocytes – predominantly included cells from FHP or IPF patients (p<2.2e-16) **(Figure 2D-G)**. CCL3 and CCL4 recruit myeloid and T cells to inflammatory sites and are increased in tissues and peripheral blood of patients with IPF and systemic sclerosis [27, 28]. S100A family proteins – including S100A8, S100A9, and S100A12 – are associated with disease presence and severity in patients with IPF [29]. Meanwhile, S100A^hi^ monocytes are associated with worse outcomes in COVID-19 [26].

Pathway analyses of top differentially expressed genes was done in EnrichR [30]. Genes enriched in the CCL3^hi^/CCL4^hi^, fibrosis-enriched monocyte subtype include IL-18 signaling pathway (*IER3, CXCL3, NFKBIZ, TNFAIP3, NFKBIA, TNF, CXCL8, IL1B, ICAM1, CCL3, CCL4, CXCL2;* p=8.9e-17; WikiPathway 2021), lung fibrosis (*TNF, CXCL8, IL1B, CCL3, CCL4, CXCL2;* p=1.5e-8; WikiPathway 2021), TNFα signaling via NFκB (*DUPS2, IER3, CXCL2, CD83, CXCL3, G0S2, TNFAIP3, PPP1R15A, TNF, IL1B, CCL4, NFKBIA, ICAM1, PDE4B*; p=1.3e-25; MSigDB Hallmark 2020), and IL-10 signaling (*IL1R2, ICAM1, IL1RN, IL1B, TNF, CCL3L1, CCL3, CCL4, CXCL2, CXCL8;* p=5.4e-18; Reactome 2022). Other genes that were also enriched in this group of cells include *SAMSN1, CD163, RNF144B, NLRP3, SOD2, BCL2A1,* and *PIM3*.

S100A^hi^ monocytes had genes highly enriched in neutrophil degranulation (*CDA, MCEMP1, CKAP4, FOLR3, GCA, MNDA, SLC2A3, QPCT, RETN, S100P, SELL, CTSD, ANPEP, ITGAM, S100A12, S100A8, S100A9*; p=3.7e-16; Reactome 2022), TLR4 cascade (*ITGAM, S100A12, S100A8, S100A9, JUN, FOS;* p=0.000077; Reactome 2022), MyD88:MAL(TIRAP) cascade initiated on plasma membrane (*CTSD, ANPEP;* p=0.00034; Reactome 2022), epithelial-to-mesenchymal transition (*JUN, BASP1, VCAN, CAPG;* p=0.0020; MSigDB Hallmark 2020), and IL-4 regulation of apoptosis (*RGS2, VCAN, JUN, ITGAM, ALOX5AP, CYP1B1, S100P, FOS*; p=0.000048; BioPlanet 2019).

To gain more insight into transcription factors that may be driving these gene expression changes noted above, we imputed transcription factor activities for monocytes. After imputing transcription factor activities, monocytes were re-clustered based on their transcription factor (TF) profiles, resulting in eight DoRothEA-based clusters **(Figure 2H)**. Chi-Squared tests showed no significant differences in regulon compositions between FHP, IPF, and controls. However, IPF and FHP had more similar regulon compositions than controls, albeit not significantly different. Clusters 4 and 6 were relatively expanded in FHP and IPF compared to controls. Cluster 5 was significantly expanded in IPF and FHP versus controls (p=0.040; p=0.024). A heatmap incorporating top variable imputed TFs revealed a “shared fibrosis signature” enriched in cluster 6 **(Figure 2I)**. EnrichR analysis identified key global regulators from the ENCODE/ChEA Consensus TF Pathways database: Ets1 (p=0.0065) and Spi1 (p=0.0064), both ETS family members known for their roles in development, cellular processes, and immune response.

### S100A^hi^ CD14+ Monocytes May Be Progenitors for SPP1+ MERTK+ Fibrosis-Associated Lung Macrophages

Myeloid cells from peripheral blood and bronchoalveolar lavage (BAL) samples were combined and re-processed, resulting in a total of 183,305 cells (123,361 PBMC and 59,944 BAL) **(Figure 3A)**. Genes of interest – including *CSF1R*, *CSF3R*, *CXCR1*, and *CCR2* – were expressed in monocytes, with *CCR2* being highly expressed in CD14+ monocytes. Fibrosis-associated CD14+ monocyte subtypes were re-identified, with one group of cells enriched in *S100A8*, *S100A9*, and *CXCL8,* and another group enriched in *ICAM1, NFKBIA, IL1B, CCL3, CCL4,* and *CCL5*. Macrophages demonstrated broad expression of genes such as *MARCO*, *CXCL3, MRC1, CCL18, CD74,* and *CSF1*, with varying expression levels in different macrophage subsets. *CSF1*, *CCL18*, *MRC1*, *SPP1, PLA2G7, MERTK, LGMN, RNASE1, CCL2,* and *CCR5* were most highly expressed in fibrosis-associated monocyte-derived macrophages (MoDMs), while *FABP4* was highly expressed in a different subset of alveolar macrophages (**Figure E6)**.

**Figure 3.**
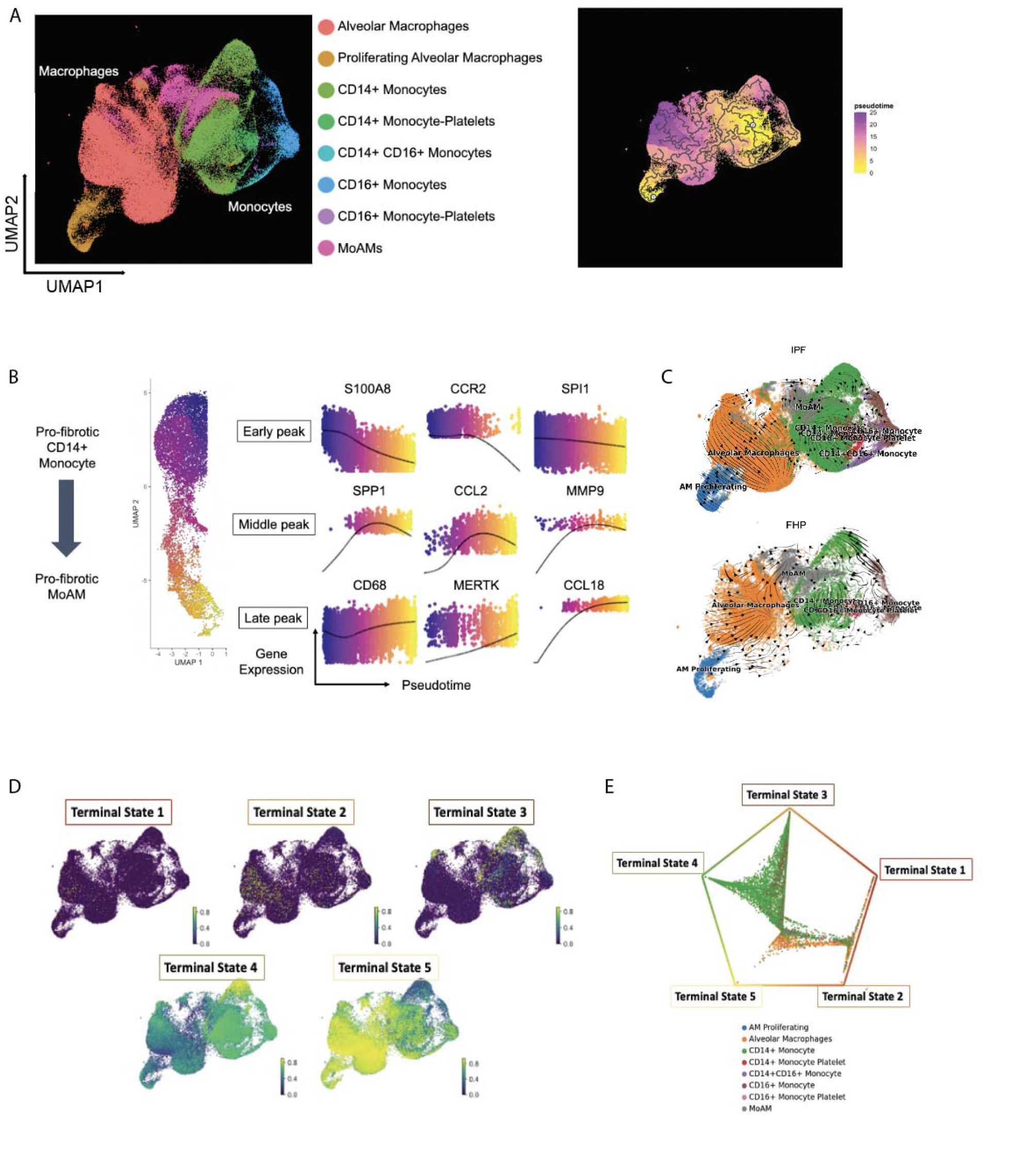
Trajectory Analysis of PBMC and BAL Myeloid Cells. **(A)** UMAP representation of 183,305 cells by cell types. **(B)** Gene expression changes over pseudotime between S100Ahi CD14+ monocyte to pro-fibrotic MoDMs, colored by pseudotime. **(C)** RNA velocity vector fields as streamlines projected onto the UMAP embedding, split by IPF and FHP. **(D)** Relative distribution of cells in the five imputed terminal states in the combined monocyte-macrophage data. **(E)** Circular embedding showing the absorption probabilities for each of the terminal states, colored by cell type annotation. PBMC: peripheral blood mononuclear cell; BAL: bronchoalveolar lavage; UMAP: uniform manifold approximation and projection; MoDM: monocyte-derived macrophage; IPF: idiopathic pulmonary fibrosis; FHP: fibrotic hypersensitivity pneumonitis.

PBMC CD14+ monocytes and BAL MoDMs were further analyzed. The trajectory root was defined based on the furthest cell from MoDMs expressing *S100A8, S100A9,* and *CD14. CCR2, SPI1, CD14,* and *S100A8* are expressed earlier in CD14+ monocytes and decrease in expression over pseudotime. The transition from monocytes to MoDMs includes a rise and fall in *CCR5*, *CEBPB*, *CCL2*, and *CSF1R* expression. An increase and subsequent decrease in *MMP9*, *CSF1*, and *SPP1* followed. Genes that continuously increase over pseudotime were *CCL18*, *CD68, MERTK, PLA2G7*, *CCL3,* and *ICAM1* (**Figure 3B)**.

RNA velocity analysis showed higher transcriptional activity in CD14+ and CD16+ monocytes (65-67% un-spliced read proportions) compared to macrophages (57-61%) [31] **(Figure E7)**. RNA velocity trajectories demonstrated that IPF and FHP monocytes differentiate into MoDMs rather than directly into alveolar macrophages **(Figure 3C)**. CD14+ monocytes also became CD16+ monocytes through intermediate monocytes.

CellRank was used to analyze combined IPF and FHP samples [32]. Five terminal states were identified, mainly CD14+ monocytes or alveolar macrophages **(Figure 3D)**. Most monocytes’ trajectories resulted in terminal states 3 or 4, corresponding to MoDMs (terminal state 3) and monocytes and alveolar macrophages (terminal state 4). Meanwhile, cells starting as alveolar macrophages remained those cells (terminal states 1, 2, and 5).

### Single-Cell Reveals Emergence of FHP-Enriched GZMK^hi^ CD8 T Cell Subpopulation

T lymphocytes were re-processed, resulting in 144,461 cells and eleven annotated subtypes **(Figure 4A-B)**. GZMK^hi^ versus GZMH^hi^ CD8+ T lymphocytes were classified based on molecular markers previously published by Perez *et al.* [33] **(Figure E8)**. GZMK^hi^ CD8+ T cells were enriched in FHP patients (p=0.0002) compared to non-FHP, IPF, and healthy controls **(Figure 4C-E)**. Enriched genes in these cells were associated with TNFα signaling via NFκB signaling (*CCL5, NFKBIA, CD68, CCL4, JUN, GADD45B, ID2, DUSP1, DUSP2, TNFAIP3, ZFP36;* p=3.4e-14; MSigDB Hallmark 2020), IL-2 signaling pathway (*ZFP36, JUN, DUSP1, CCL4, CCL5, NKG7, CXCR4, GZMA, DUSP6, CTSW, GZMK, CD69*; p=7.7e-9; BioPlanet 2019), IL-1 signaling pathway regulation of extracellular matrix (*ZFP36, JUN, TNFAIP3, NFKBIA, CCL4, CCL5, DUSP6*; p=3.2e-8; BioPlanet 2019), and IL-12-mediated signaling events (*GZMA, GADD45B, CCL4L2, CD8A, CD8B, HLA-DRB1;* p=3.2e-8; BioPlanet 2019).

**Figure 4.**
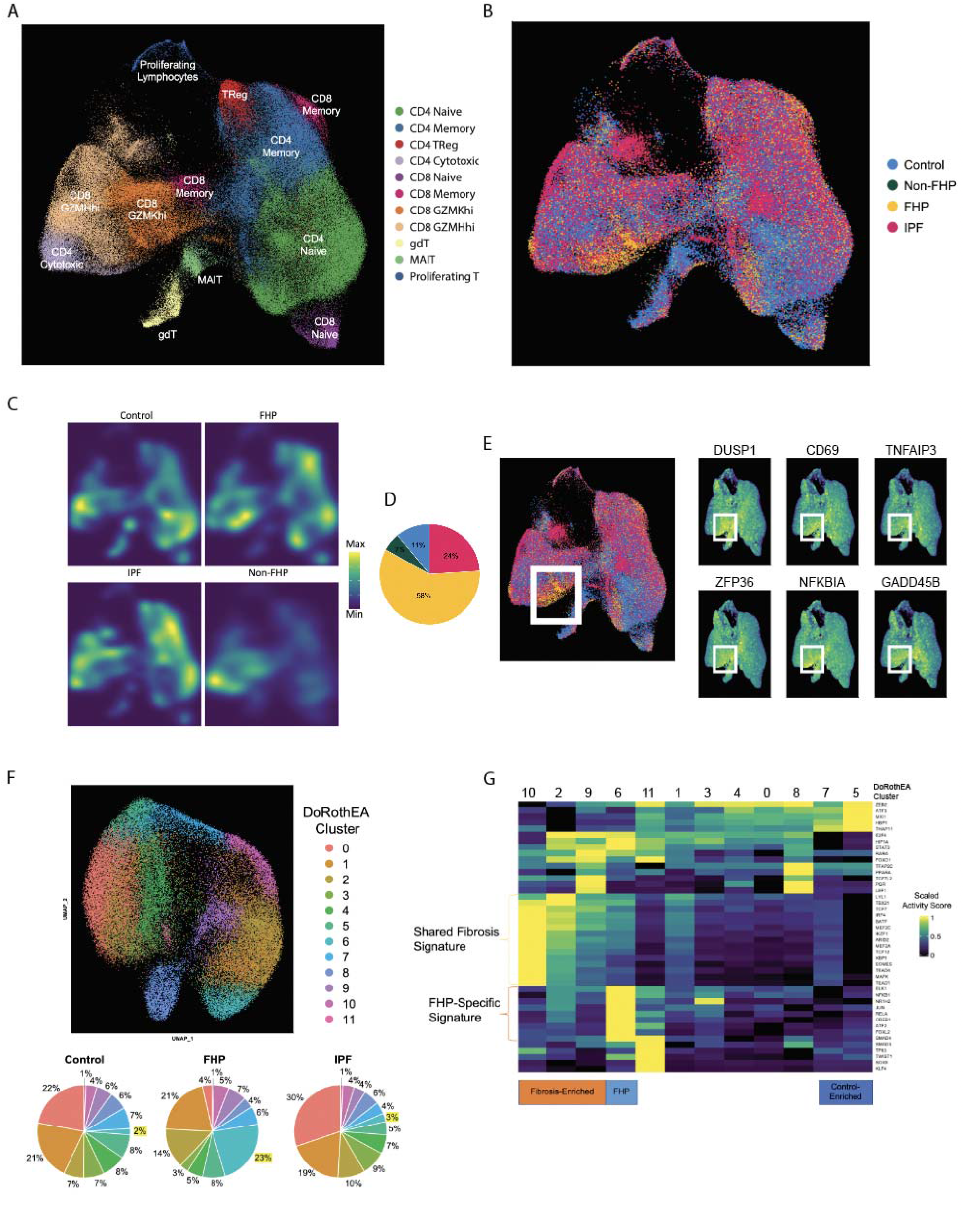
CD8+ T Cell Subtypes Enriched in FHP. UMAP representation of 144,461 cells by **(A)** T cell subtypes and **(B)** disease subtypes**. (C)** Density graph by disease subtype. **(D)** Breakdown of FHP-enriched GZMKhi CD8+ T cells (p=0.0002) by disease. **(E)** Gene expression of *DUSP1, CD69, TNFAIP3, ZFP36, NFKBIA,* and *GADD45B.* **(F)** UMAP of CD8+ T cells clustered by DoRothEA-VIPER transcription factor activity scores, and the breakdown of the CD8 T lymphocyte DoRothEA clusters by control, FHP, and IPF. **(G)** Heatmap showing transcription factors with most variable imputed activities across CD8+ T cell DoRothEA clusters. UMAP: uniform manifold approximation and projection; IPF: idiopathic pulmonar fibrosis; FHP: fibrotic hypersensitivity pneumonitis; TF: transcription factor.

CD8 T lymphocytes were re-clustered based on TF activity scores, yielding twelve DoRothEA clusters **(Figure 4F)**. The compositions of DoRothEA clusters differed significantly in FHP versus IPF and controls (p=1.1e-5; p=1.4e-4) but not between IPF and controls (p=0.9541). Cluster 0 was decreased in FHP compared to IPF and controls (p=8.5e-7; p=2.2e-4), while cluster 6 was increased versus IPF and controls (p=2.9e-5; p=5.8e-6). Clusters 2, 9, and 10 were expanded in FHP and IPF and shared transcription factors enriched in Gata2 activity (p=0.025; ChEA 2022). Cluster 6, specific to FHP, showed enriched transcription factors, including *ELK1*, *NFKB1, RELA, JUN,* and *SMAD4*. *ELK1* regulates cell proliferation and survival and regulates AVB6, activating TGFβ [34] **(Figure 4G)**. Though no significant enrichments were found in ChEA and ENCODE TF databases, the transcription factors in cluster 6 were enriched in pathways such as TGFβ signaling (p=4.3e-6; Elsevier Pathway Collection) and TNFα signaling via NFκB (p=7.1e-4; MSigDB Hallmark 2022).

### Predominantly FHP Patients Demonstrate Activated B Cell Phenotype

B lymphocytes and plasma cells were re-clustered, resulting in 28,108 cells and four distinct cell types: naïve B cells, memory B cells, intermediate B cells, and plasma cells. Isoform expression patterns concorded with the literature [35] (**Figure 5A-C; Figure E9)**. Namely, plasma cells, memory B cells, and intermediate B cells exhibited higher expression of IgA and IgG isoforms, whereas IgM and IgD heavy chains were more expressed in naïve B cells. IgE expression was relatively low, primarily observed in memory B cells **(Figure E9)**. A subset of naïve B cells was predominantly expanded in FHP (p=5.7e-10) **(Figure 5D-E)**. These cells showed enrichment in genes involved in signaling by NGF and antigen-activated B cell receptor generation of second messengers (*FOXO1, BRAF, AKT3, PRKCE, ITPR1, NFKB1, VAV3, LYN, TRIO;* p=4.6e-7; BioPlanet 2019), and B cell receptor signaling pathway (*LYN, BRAF, NFKB1, FOXO1;* p=0.0011; WikiPathway 2021).

**Figure 5.**
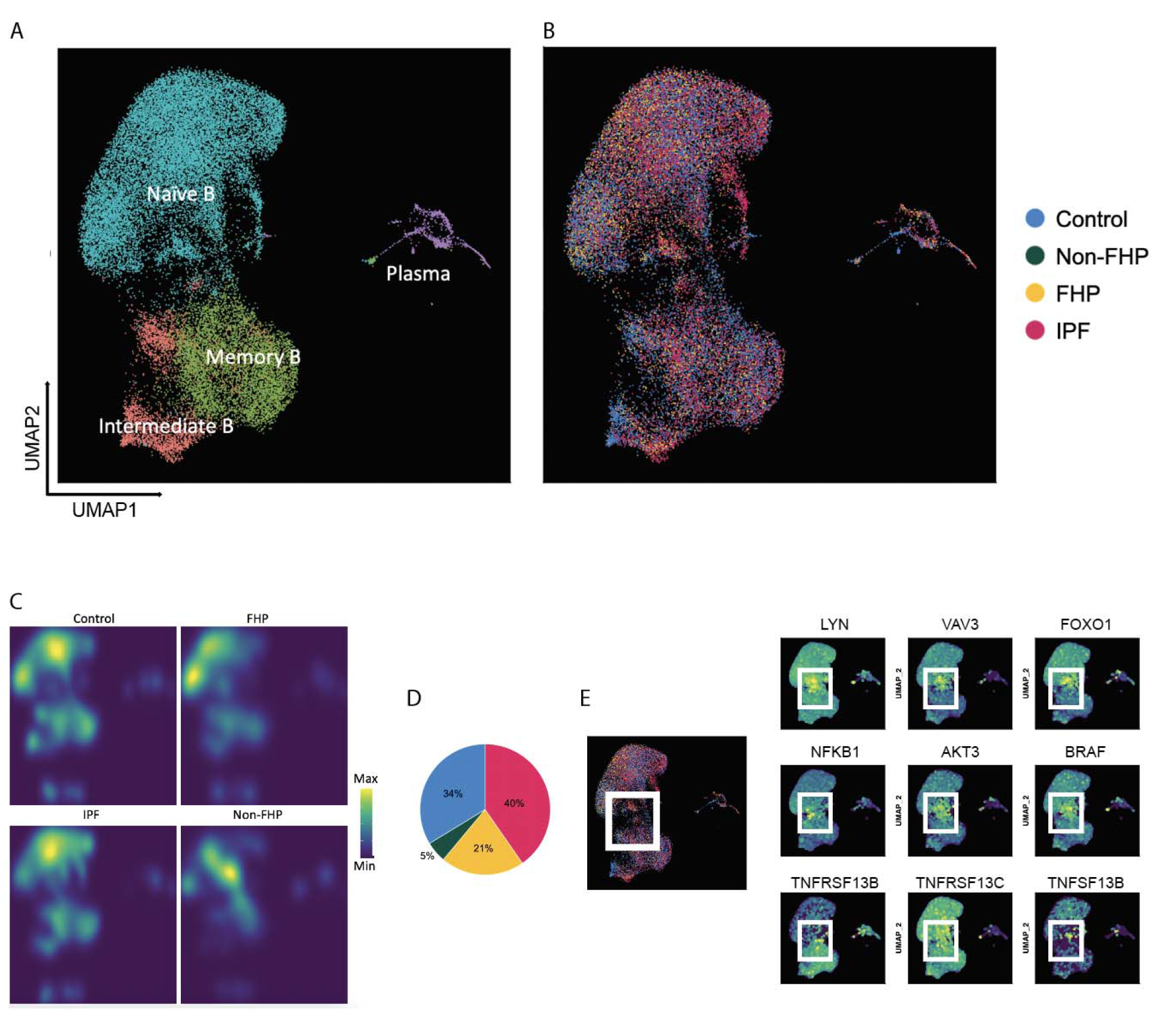
B cell subpopulations enriched in FHP. UMAP representation of 28,108 B cells by **(A)** cell subtypes and **(B)** disease subtypes. **(C)** Density graph of B cells by disease subtype. **(D)** Breakdown of FHP-enriched naïve B cells (p=5.7e-10) by disease. **(E)** Gene expression of *LYN, VAV3, FOXO1, NFKB1, AKT3, BRAF, TNFRSF13B, TNFRSF13C,* and *TNFSF13B.* UMAP: uniform manifold approximation and projection; IPF: idiopathic pulmonary fibrosis; FHP: fibrotic hypersensitivity pneumonitis.

### Fibrogenic Immune Signaling Identified in Fibrotic Lung Disease Patients Versus Healthy Volunteers

Ligand-receptor pairs across different cell types were investigated to examine signaling differences between fibrotic lung disease (IPF and FHP) patients and healthy controls **(Figure 6A-D)**. In fibrotic disease, ligands such as *HBEGF* and *SPON2* were increased. Fibrotic diseases also showed increased ligand expression of *IL1B* and *IGF1R*. Comparing healthy controls and fibrotic disease, the latter displayed enrichment in IL-2/STAT5 pathways (*SELL, IL2RB, LTB, CD44, IGF1R;* p=7.1e-7) and inflammatory responses (*SELL, IL2RB, CD55, HBEGF*; p=2.3e-5; MSigDB Hallmark, 2020), while healthy controls demonstrated enrichment in IL-6/JAK/STAT3 signaling pathways (*ITGA4, IFNGR2, TNF, CD44, TLR2*, p=1.6e-7; MSigDB Hallmark, 2020). Next, distinguishing between FHP and IPF revealed FHP-enriched ligand-receptor pairings, including *CCL5* to *CXCR3*. Ligands and receptors increased in IPF include *IL1B*, *TGFB1*, *IGF2R*, *CD36*, and *ITGB3* (**Figure 6E-H)**.

**Figure 6.**
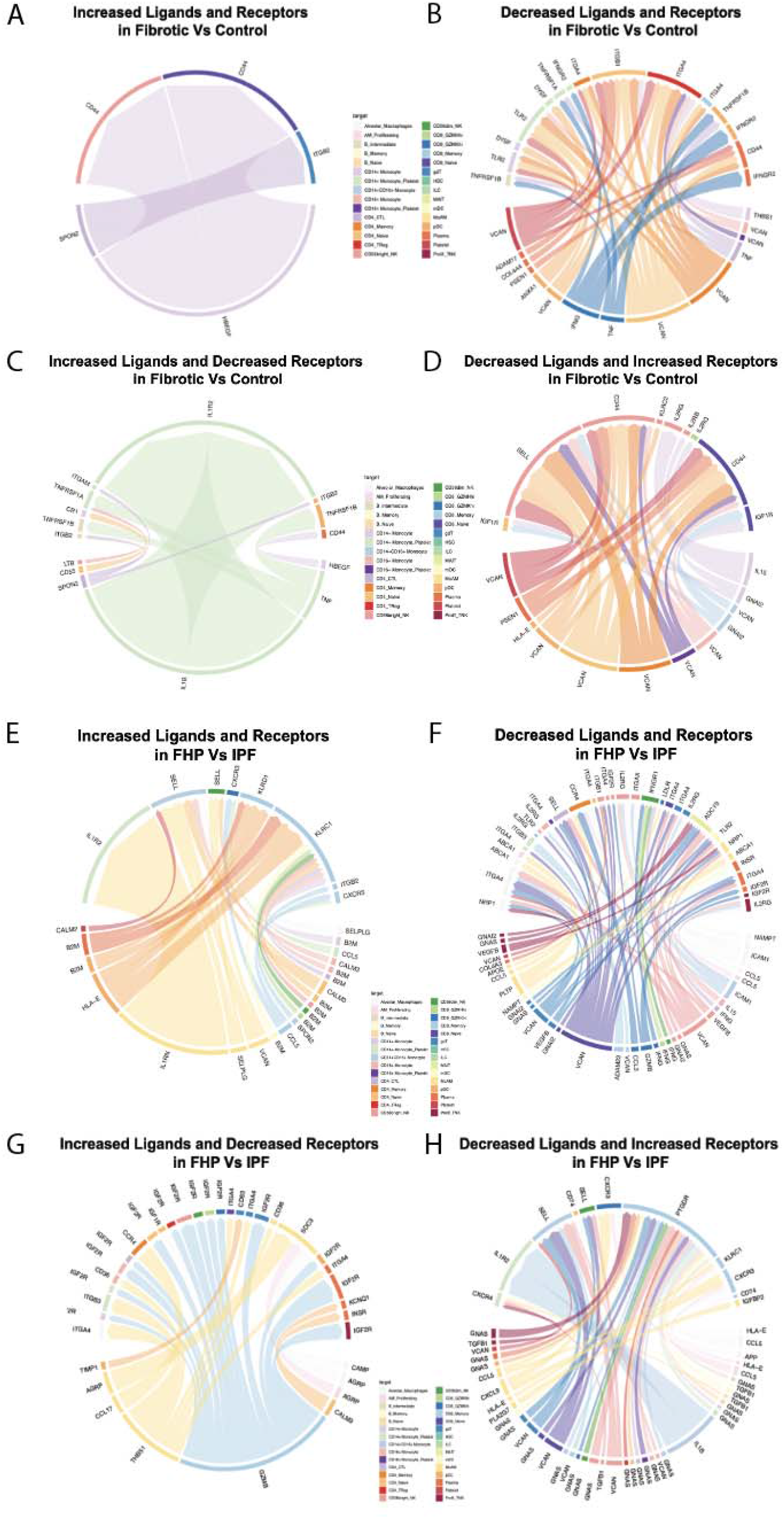
Aberrant Cell-Cell Communication in Disease. **(A-D)** Circos plots demonstrating perturbed ligand-receptor pairs in fibrotic disease versus healthy controls. The bottom half of each plot represents the ligands, while the top half represents the receptors. Sending and receiving cell types are colored by cell type. Ligands and receptors increased **(A)** and decreased **(B)** in fibrotic disease versus controls. **(C)** Increased ligands and decreased receptors in fibrotic versus control **(D)** decreased ligands and increased receptors in fibrotic versus control. **(E-H)** Circos plots demonstrating most significantly perturbed ligand-receptor pairs in FHP and IPF. Ligands and receptors increased **(E)** and decreased **(F)** in FHP versus IPF. **(G)** Increased ligands and decreased receptors in FHP versus IPF. **(H)** Decreased ligands and increased receptors in FHP versus IPF. IPF: idiopathic pulmonary fibrosis; FHP: fibrotic hypersensitivity pneumonitis.

### Immune Receptor Repertoire Analysis Reveals Differential Isotype Enrichment in IPF and FHP B Cells

To investigate T and B cell receptor (TCR, BCR) repertoires in fibrotic lung disease, single-cell V(D)J analysis was performed on IPF, FHP, and healthy control samples. 49,556 unique T cell clones and 12,008 unique B cell clones were identified **(Figure 7A-B)**. No significant differences in T and B cell receptor repertoire diversities were observed among disease groups and controls **(Figure E10)**. However, FHP patients had more distinct T and B cell complementarity-determining region 3 (CDR3) amino acid usage profiles than IPF and controls **(Figure 7C-D)**.

**Figure 7.**
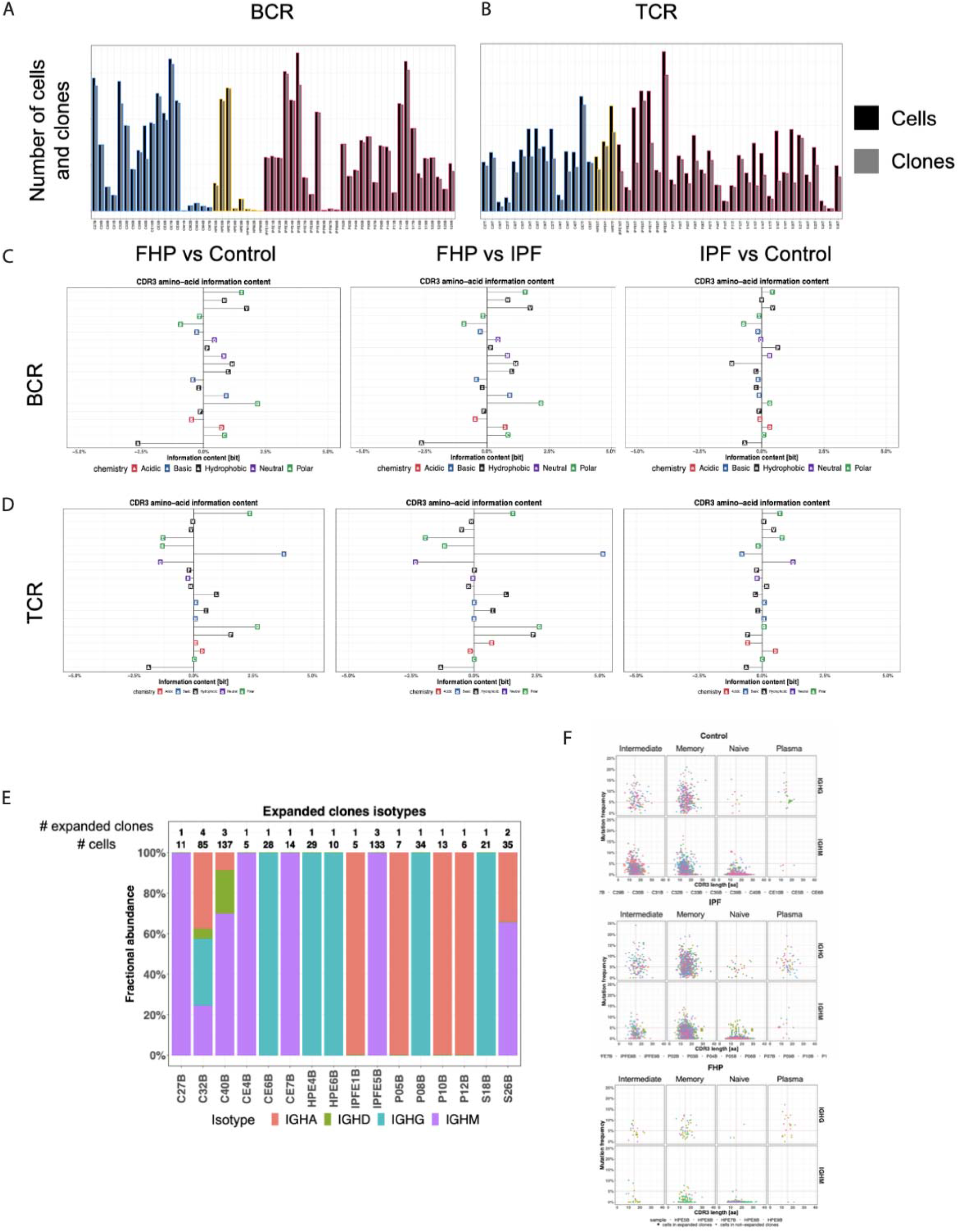
BCR and TCR Repertoire Analyses. Number of cells versus number of clones from each sample grouped by disease subtype for **(A)** BCRs and **(B)** TCRs. **(C)** BCR and **(D)** TCR CDR3 amino acid usage as percent change of information content for FHP patients relative to control; FHP patients relative to IPF; and IPF patients versus control, colored by the amino acid chemistry (acidic, basic, hydrophobic, neutral, and polar). **(E)** Fractional abundance of expanded B cell clones’ isotypes per sample with expanded clones. **(F)** Mutation frequency plotted against CDR3 amino acid length and mutation frequency for healthy controls, IPF, and FHP, split by B cell subtypes and BCR isotype. The horizontal point represents 5% mutational frequency while the vertical line represents an amino acid length of 15 residues. Cells in expanded clones are denoted by larger point size. BCR: B cell receptor; TCR: T cell receptor; CDR3: complementarity determining region 3; IPF: idiopathic pulmonary fibrosis; FHP: fibrotic hypersensitivity pneumonitis.

Further analysis focused on expanded B cell clones, which demonstrated significant differences in isotype representation **(Figure 7E)**. B cell clones from FHP patients were more enriched in IgG isotypes compared to IPF and controls (p<2.2e-16). IPF clones showed higher IgM and IgA isotype enrichment compared to FHP (p=2.4e-15; p=0.0025) and controls (p=5.2e-5; p=0.052), respectively. Notably, memory B and plasma cells exhibited higher IgG mutation rates (>5%) compared to IgM mutation rates, indicating ongoing somatic hypermutation in these cell subsets (p<0.05). Analysis of CDR3 regions revealed specific subtypes with significantly longer amino acid residues (p<0.05), including memory B (IgG and IgM), naïve B (IgM), and intermediate B (IgG and IgM) in IPF; naïve B cells (IgM) in FHP; and memory B (IgG), naive B (IgM), and plasma cells (IgG) in controls **(Figure 7E).**

## Discussion

In this study, we conducted comprehensive single-cell RNA-sequencing of PBMC and BAL samples from patients with IPF, FHP, and healthy controls. Our findings revealed shared and disease-specific alternations in immune cell populations and gene expression profiles, shedding light on the underlying immune mechanisms of these fibrotic lung diseases. Our study also represents the first combined peripheral blood single-cell atlas of IPF and FHP to date.

In terms of cell composition, both IPF and FHP patients exhibited increased monocyte proportions and decreased T and B cell subpopulations in peripheral blood compared to controls. IPF patients showed elevated platelet-monocyte complexes, suggesting platelet activation specific to this disease. FHP patients had higher platelet counts, indicating potential platelet involvement in disease pathogenesis. These findings align with previous reports of elevated CD14+ monocytes in fibrotic conditions and activated platelets and increased platelets in IPF and FHP, respectively [12, 16, 18, 36–38]. IPF patients displayed an age-independent decrease in naïve T cells and a relative increase in memory T cells, suggesting prior antigen sensitization [39]. FHP and IPF patients exhibited reduced counts of MAIT cells, possibly indicating their migration to the lungs and contribution to the pro-inflammatory and pro-fibrotic environment, like the role of MAIT cells in COVID-19 and COPD [40, 41]. Additionally, Tγδ cells were decreased in FHP patients’ BAL fluid, suggesting possible migration to lung tissues [42, 43]. CD56bright NK cells were reduced in FHP compared to controls and IPF, consistent with systemic inflammation-associated diseases [44–47]. Within B cells, FHP patients showed increased naïve B cells and decreased memory B cells, resembling dysregulation seen in rheumatoid arthritis-ILD [48, 49]. These findings suggest that B cell dysregulation and disease pathogenesis in FHP may more closely resemble connective tissue disorder associated ILDs than IPF.

Moreover, we analyzed gene expression profiles of myeloid and lymphoid populations across diseases. Both IPF and FHP showed enrichment of pro-fibrotic CCL3/CCL4^hi^ and S100A^hi^ CD14+ monocyte subpopulations, while FHP exhibited unique gene expression signatures in CD8+ T and B cells, characterized by pro-lymphocyte activation and pro-inflammatory profiles. BAL CCL3 and CCL4 expression are strongly associated with mortality in IPF [50]. S100A proteins activate TLR4 and the downstream inflammatory cascade. They are increased in both blood and BAL fluid of IPF patients and associated with worse pulmonary function tests [29]. High S100A expression in PBMCs is also linked with worse COVID-19 outcomes [26]. By integrating PBMC and BAL samples, we found evidence suggesting differentiation of S100A^hi^ CD14+ monocytes into SPP1^hi^ macrophages in the lungs. Pseudotime and RNA velocity analyses supported this differentiation trajectory, highlighting the dynamic gene expression patterns during disease progression. Previous research by Adams *et al.* demonstrated early expression of *SPP1* and *LIPA*, followed by later expression *SPARC, PALLD, GPC4, CTSK,* and *MMP9* in the IPF-associated macrophage archetype [19, 51]. In our study, fibrotic MoDM-associated genes – *MMP9, SPP1, CD68, CCL18, MERTK,* and *PLA2G7* – were expressed in MoDMs, but did not all express contemporaneously, suggesting a much more dynamical role of these fibrosis-associated cell types.

The activated GZMK^hi^ CD8+ T cells, which were enriched in FHP, have been observed in various autoimmune diseases. For instance, Jonsson *et al.* identified highly activated *GZMK-*expressing CD8+ T cells in synovial tissues of rheumatoid arthritis patients, as well as in inflamed tissues of other autoimmune diseases and COVID-19 [52]. Xu *et al.* described an expanded GZMK+ CXCR6+ CD8+ T cell subset in patients with primary Sjogren’s syndrome, suggesting their role as precursors of tissue-resident T cells [53]. In FHP, Wang *et al*. demonstrated the expansion of effector CD8+ T cells in the lung, potentially originating from the GZMK+ CD8+ T cells [54]. Notably, *GZMK* expression in CD8+ T cells has also been associated with neutrophil activity in colorectal cancer patients [55]. Although similarities exist with other autoimmune diseases, the FHP-enriched GZMK+ CD8+ T cell subpopulation has distinct characteristics. For example, in systemic lupus erythematosus (SLE) patients, CD8+ GZMHhi, and not GZMKhi, T cells were expanded [33]. Furthermore, transcription factor analyses supported the FHP-associated signatures in CD8+ T cells, suggesting that FHP may be driven by foreign and autoantigens, leading to a disease-specific adaptive immune response mediated by these T cells. In contrast, IPF lacks this specific population, indicating a potential T-cell-independent process in this disease.

Differentially expressed ligands and receptors were identified in our study. Fibrotic diseases showed increased expression of *HBEGF*, *SPON2*, *IL1B,* and *IGF1R*. *HBEGF*, predominantly expressed in myeloid lineages, promotes epithelial-to-mesenchyme transition through *MMP7* and *MMP9*, cell invasion, and angiogenesis [56–59]. *SPON2,* expressed in T cells, acts as a recruitment signal for inflammatory cells [60, 61]. *IL1B* contributes to extracellular matrix production mediated by Smad2/3, NFκB, and TGFβ pathway, and *IGF1R* is linked to fibrogenesis [62–67]. FHP exhibited specific enrichment of CXCR3 in T cells, which is associated with autoimmune diseases, including multiple sclerosis, ulcerative colitis, and rheumatoid arthritis [68]. These findings indicate dysregulation of cell-cell signaling in fibrosis-associated pathways, as well as more enhanced T cell signaling in FHP.

Finally, the immune receptor repertoires were explored. FHP had greater differences in T and B cell CDR3 region amino acid composition than IPF and healthy controls. Notably, expanded B cell clones were significantly skewed towards IgA and IgM isotypes for IPF and IgG for FHP, suggesting distinct adaptive immune responses in disease. These findings align with previous studies showing increased IgM- and IgA-secreting cells in healthy B cells co-cultured with IPF PBMC supernatants [69], and the association of IgGs with serum antibodies in fibrotic lung diseases [52, 70–75]. The IgA predominance in IPF may be due to TGFβ-induced class switching in a fibrotic environment [76]. These results suggest an earlier and mucosal-based B cell-mediated response in IPF, while FHP exhibits a more mature or chronic immune response. Further validation is required to identify the specific antigens driving IPF and FHP, if any.

This study had several limitations. Since there were only five non-FHP patients, there were few statistically significant differences in cell composition and gene expression in non-FHP versus other disease and control groups. Interestingly, CD4+ cytotoxic T cells were increased solely in non-FHP compared to controls, suggesting an inflammatory response akin to those seen in viral and autoimmune conditions [77], while non-FHP patients had fewer memory CD4+ T cells than those with FHP. Though the sample numbers are small, these findings may indicate that non-FHP patients launch a more robust and unprimed inflammatory response than FHP. Other limitations include a small overall cohort size that precluded ROC and more in-depth immune repertoire analyses, heterogeneous treatment, and the need for further mechanistic validations. The observed heterogeneity may stem from technical variability, genetic and epigenetic factors, and individual lifestyle differences, all influencing cellular behavior and gene expression. We tried to mitigate these factors by selecting patient cohorts with similar age, sex, ethnicities, and pulmonary function tests. Despite these limitations, our findings have aligned with current literature and are hypothesis-generating. Validation of these findings are out of the scope of this study and may be completed in a larger cohort by flow cytometry.

In conclusion, our study revealed shared fibrosis-related and disease-specific signatures in the immune cells of IPF and FHP patients, supporting the notion that, although the cellular and molecular mechanisms involved in the initiation of these diseases likely differ, during the development and progression of the fibrotic response they share some likely general profibrotic mechanisms. The goal is to improve diagnosis and treatment for patients by using easily accessible peripheral blood. Circulating S100A^hi^ and CCL3/CCL4^hi^ CD14+ monocytes can distinguish whether a patient has fibrotic lung disease, and GZMK^hi^ CD8+ T cell subtypes can determine whether a patient has FHP over IPF. Targeting pro-fibrotic monocytes and disease-associated alveolar macrophages could attenuate fibrosis. Finally, shifting towards a classification schema based on immune endotypes – whether the patient has “intrinsic” versus “extrinsic” immune signatures – may lead to more targeted therapies and improved outcomes for patients with pulmonary fibrosis.

## Notes

**Conflict of Interest:** NK is a scientific founder at Thyron, served as a consultant to Boehringer Ingelheim, Pliant, Three Lake Partners, Astra Zeneca, RohBar, Veracyte, Augmanity, CSL Behring, Splisense, Galapagos, Fibrogen, GSK, Merck and Thyron over the last 3 years, reports Equity in Pliant and Thyron, and grants from Veracyte, Boehringer Ingelheim, BMS and non-financial support from Astra Zeneca. AU reports receiving research funding from Boehringer Ingelheim, and personal consulting fees or honoraria from Boehringer Ingelheim, Kamada, RemedyCell, Augmanity Nano, Splisense, Veracyte, and 1E Therapeutics in the last 3 years. JCS reports receiving honoraria and travel support from Boehringer Ingelheim and Kinevant. AJ reports support from Fond de dotation du Souffle, Philippe Foundation, Bourse de Mobilite CHU de Caen, Bourse de mobilite internation interregion Nord Ouest G4. AP reports honoraria and travel expenses from Boehringer Ingelheim, Novartis, GILEAD, Nitto-Denko, and CSL Behring. All other authors report no conflict of interest.

### Competing Interest Statement

NK is a scientific founder at Thyron, served as a consultant to Boehringer Ingelheim, Pliant, Three Lake Partners, Astra Zeneca, RohBar, Veracyte, Augmanity, CSL Behring, Splisense, Galapagos, Fibrogen, GSK, Merck and Thyron over the last 3 years, reports Equity in Pliant and Thyron, and grants from Veracyte, Boehringer Ingelheim, BMS and non-financial support from Astra Zeneca. AU reports receiving research funding from Boehringer Ingelheim, and personal consulting fees or honoraria from Boehringer Ingelheim, Kamada, RemedyCell, Augmanity Nano, Splisense, Veracyte, and 1E Therapeutics in the last 3 years. JCS reports receiving honoraria and travel support from Boehringer Ingelheim and Kinevant. AJ reports support from Fond de dotation du Souffle, Philippe Foundation, Bourse de Mobilite CHU de Caen, Bourse de mobilite internation interregion Nord Ouest G4. AP reports honoraria and travel expenses from Boehringer Ingelheim, Novartis, GILEAD, Nitto-Denko, and CSL Behring. All other authors report no conflict of interest.

