## Supplement for "Peripheral Blood Single-Cell Sequencing Uncovers Common and Specific Immune Aberrations in Fibrotic Lung Diseases"

**DATA SUPPLEMENT**

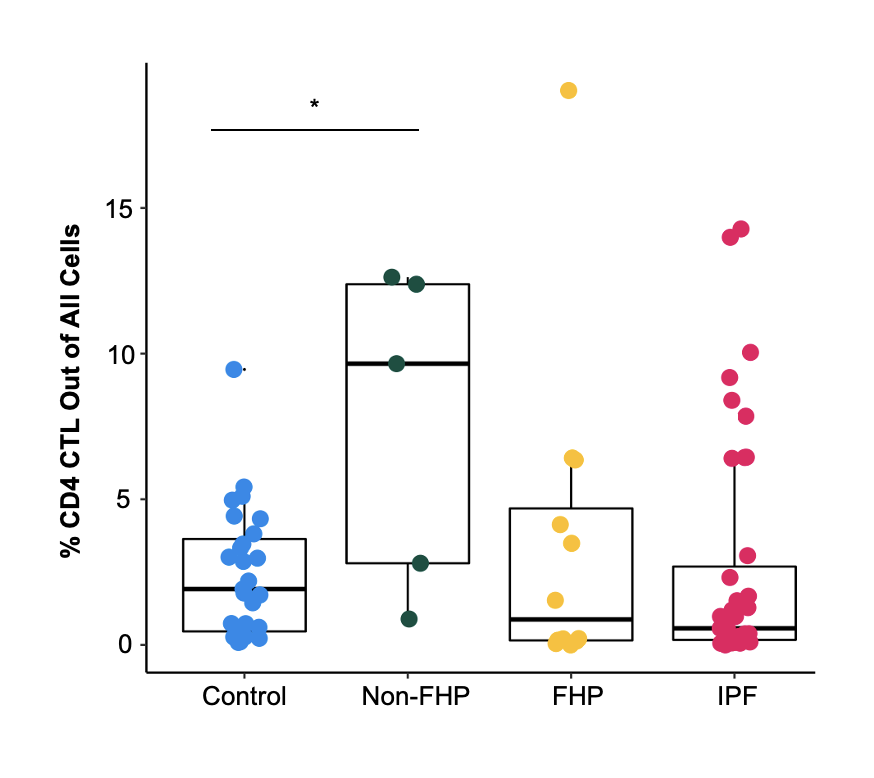

**Figure E1.** Percentage of CD4 cytotoxic T cells for each sample, grouped by disease subtype. *p<=0.05, **p<=0.01, ***p<=0.001. FHP: fibrotic hypersensitivity pneumonitis; IPF: idiopathic pulmonary fibrosis; CTL: cytotoxic T lymphocyte.

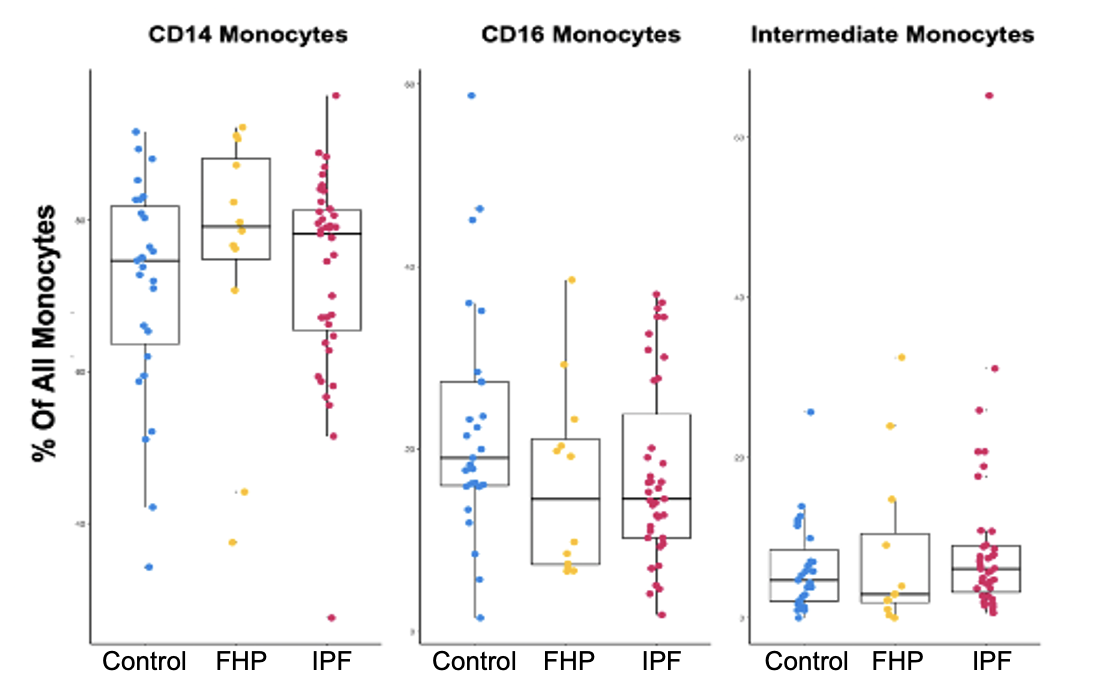

**Figure E2.** Percentage of monocyte cell type out of all monocytes for each sample, grouped by disease subtype. *p<=0.05, **p<=0.01, ***p<=0.001. FHP: fibrotic hypersensitivity pneumonitis; IPF: idiopathic pulmonary fibrosis.

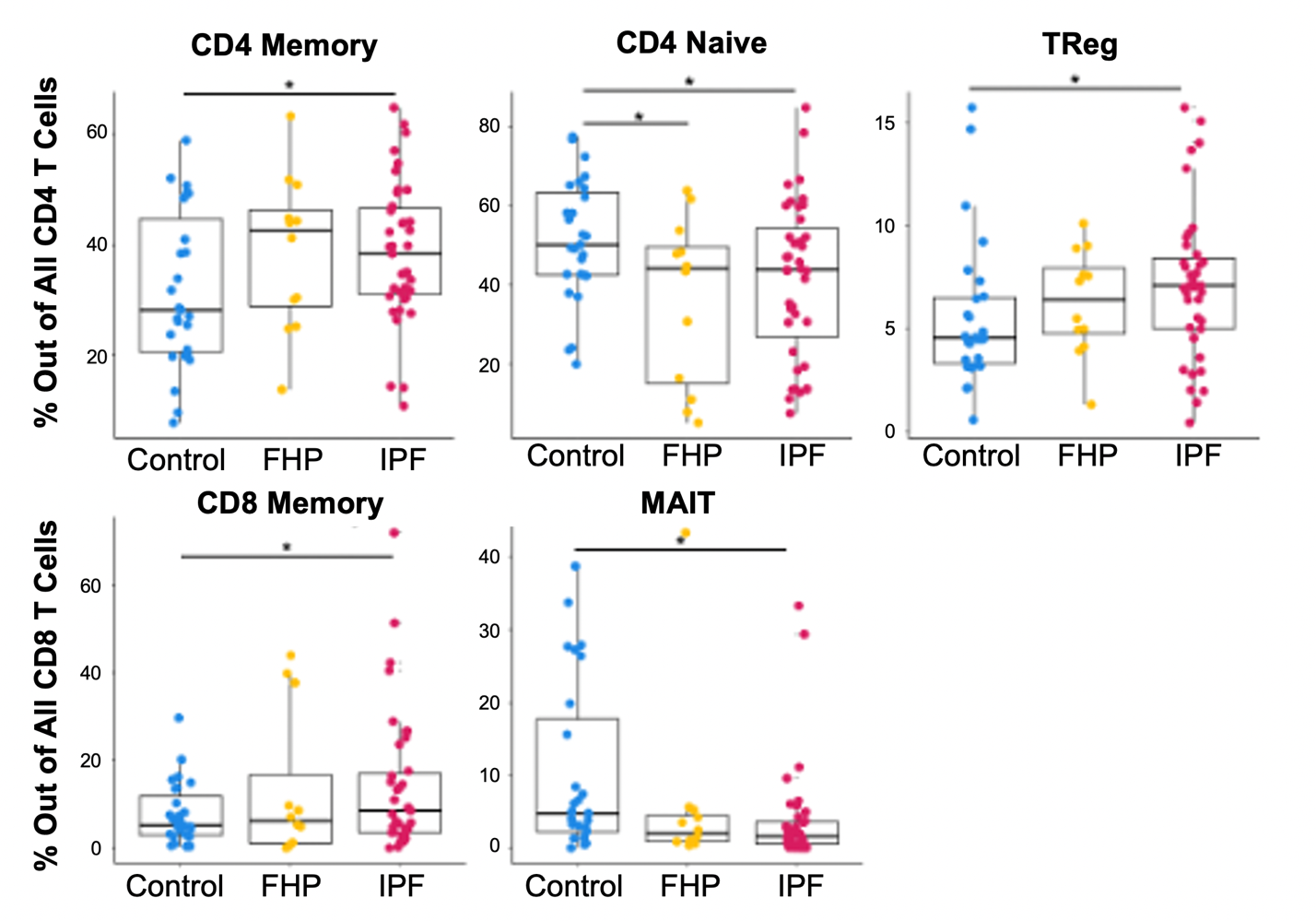

**Figure E3.** Percentage of T cell type out of (A) CD4 T cells and (B) CD8 T cells for each sample, grouped by disease subtype. *p<=0.05, **p<=0.01, ***p<=0.001. FHP: fibrotic hypersensitivity pneumonitis; IPF: idiopathic pulmonary fibrosis.

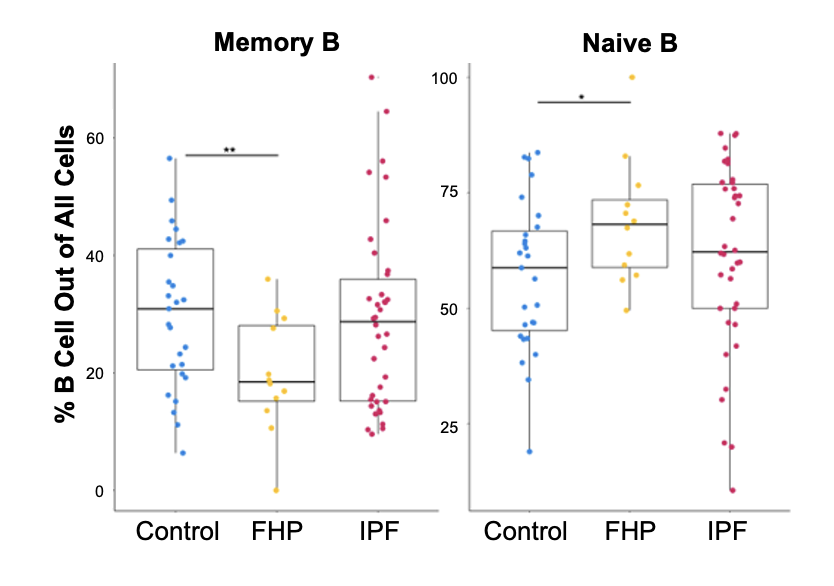

**Figure E4.** Percentage of B cell subtype out of all B cells each sample, grouped by disease subtype. *p<=0.05, **p<=0.01, ***p<=0.001. FHP: fibrotic hypersensitivity pneumonitis; IPF: idiopathic pulmonary fibrosis.

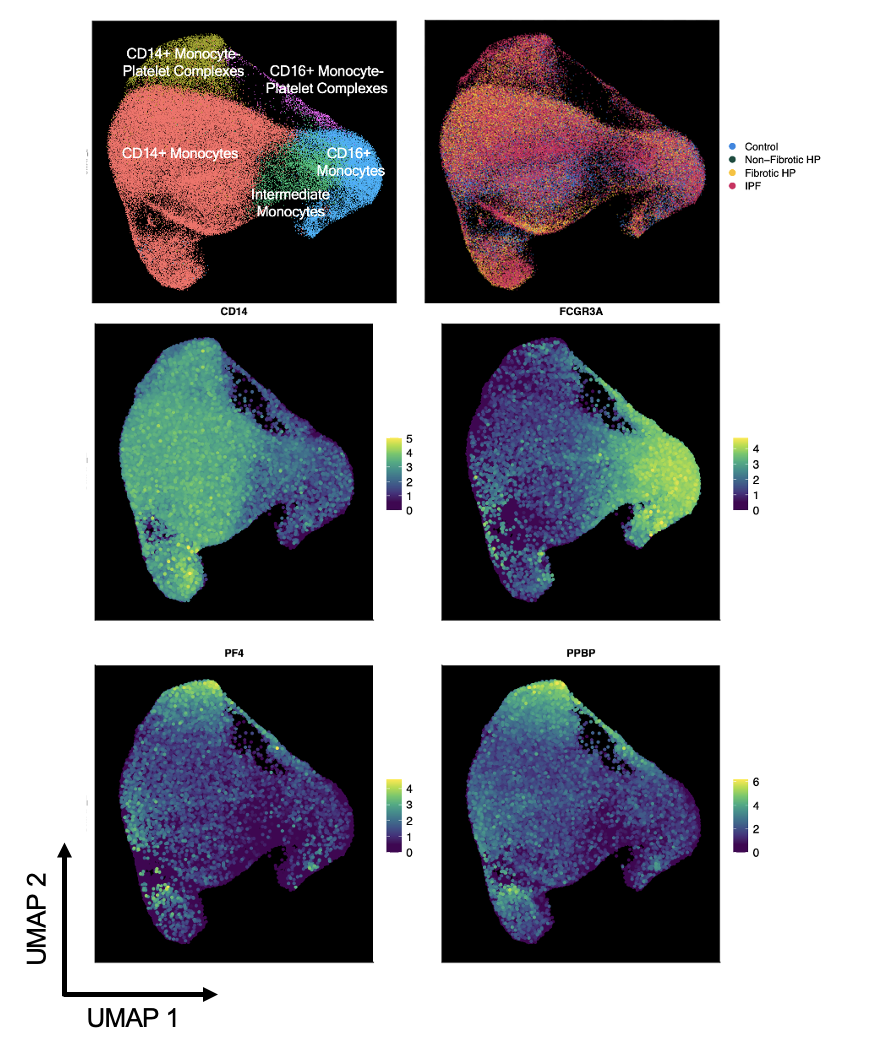

**Figure E5.** Monocyte feature plots of *CD14* (CD14+ monocytes)*, FCGR3A* (CD16+ monocytes)*, PF4/PPBP* (monocyte-platelet complexes).

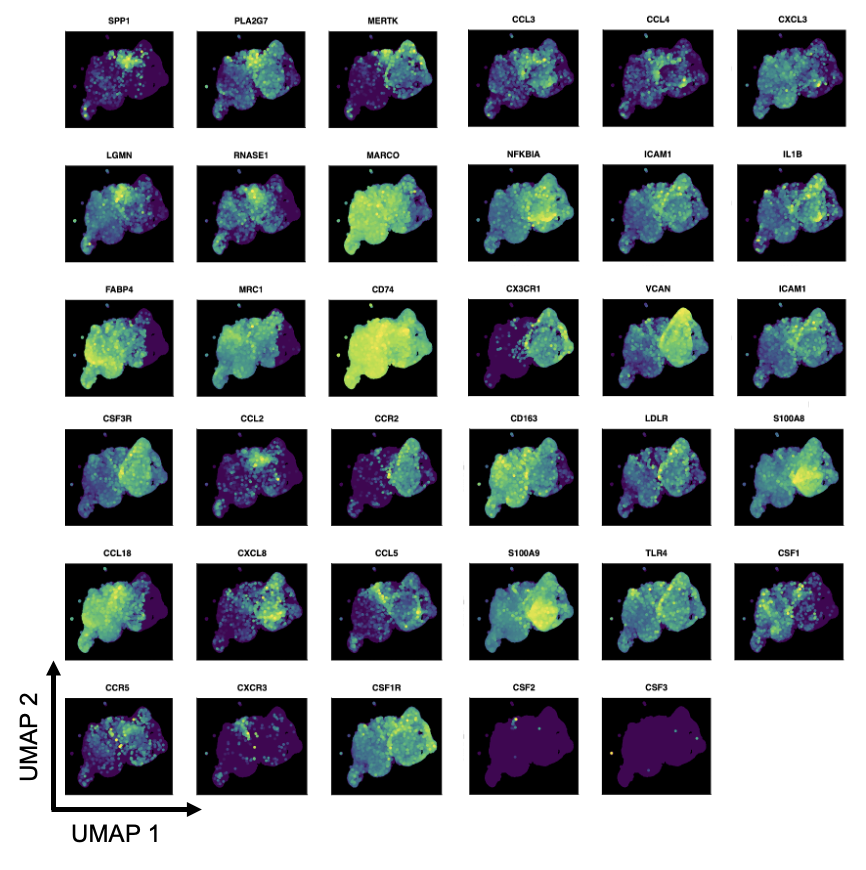

**Figure E6.** Combined BAL and PBMC monocyte and macrophage feature plots.

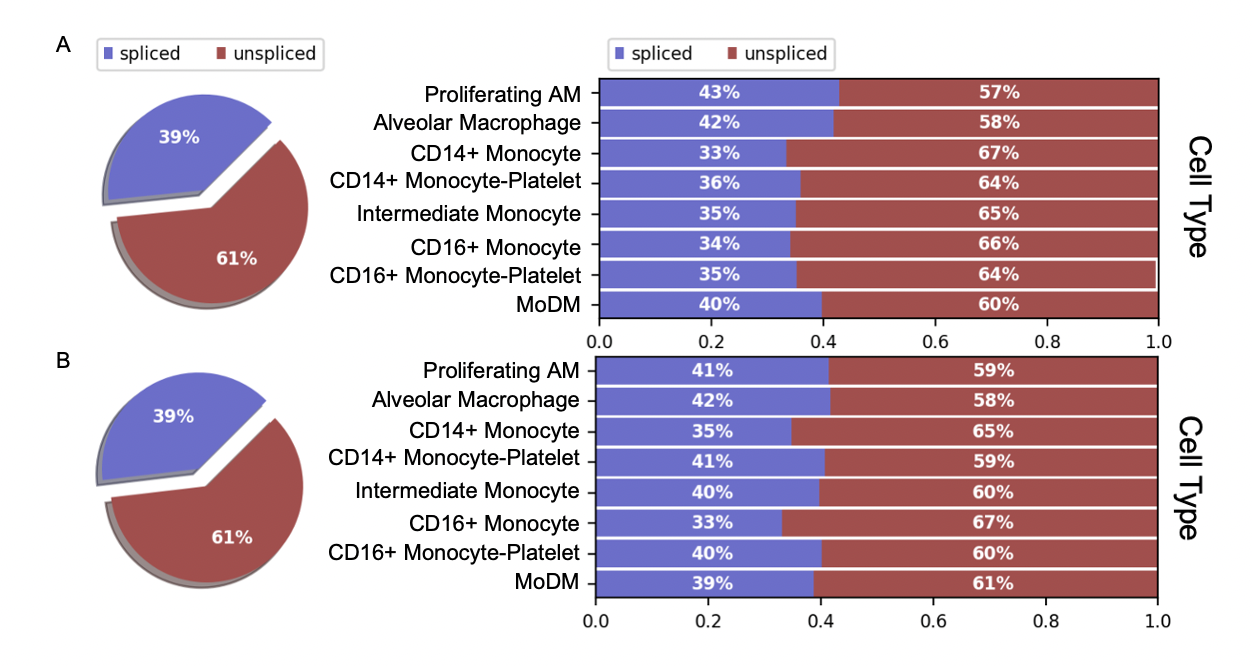

**Figure E7.** Unspliced and spliced read proportions across myeloid cell types in (A) IPF and (B) FHP combined BAL and PBMC samples. AM: alveolar macrophage; MoDM: monocyte-derived macrophage; IPF: idiopathic pulmonary fibrosis; FHP: fibrotic hypersensitivity pneumonitis.

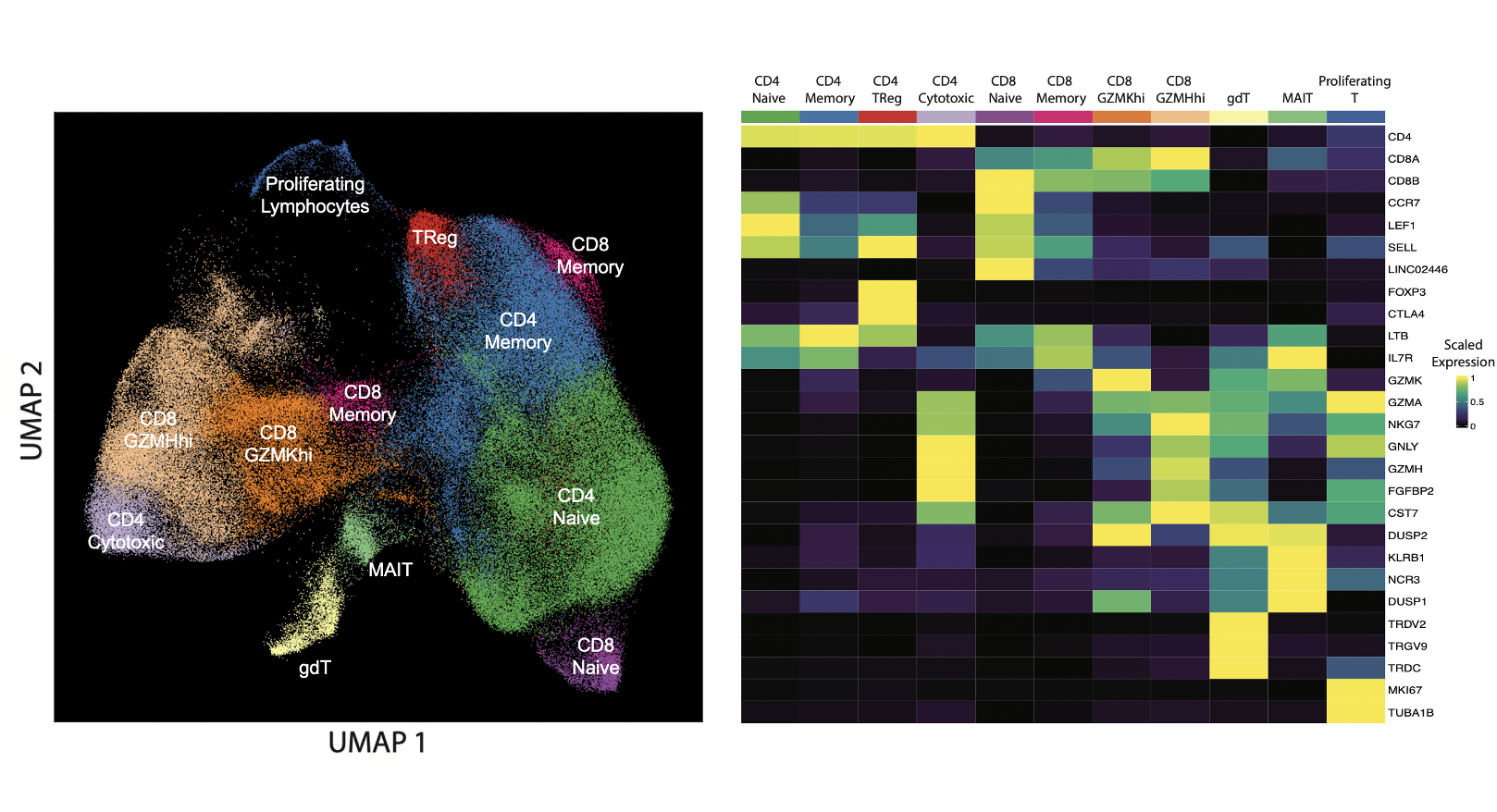

**Figure E8.** Heatmap demonstrating marker genes for each T cell subset. TReg: regulatory T cell; gdT: gamma-delta T cell; MAIT: mucosal-associated innate T cell.

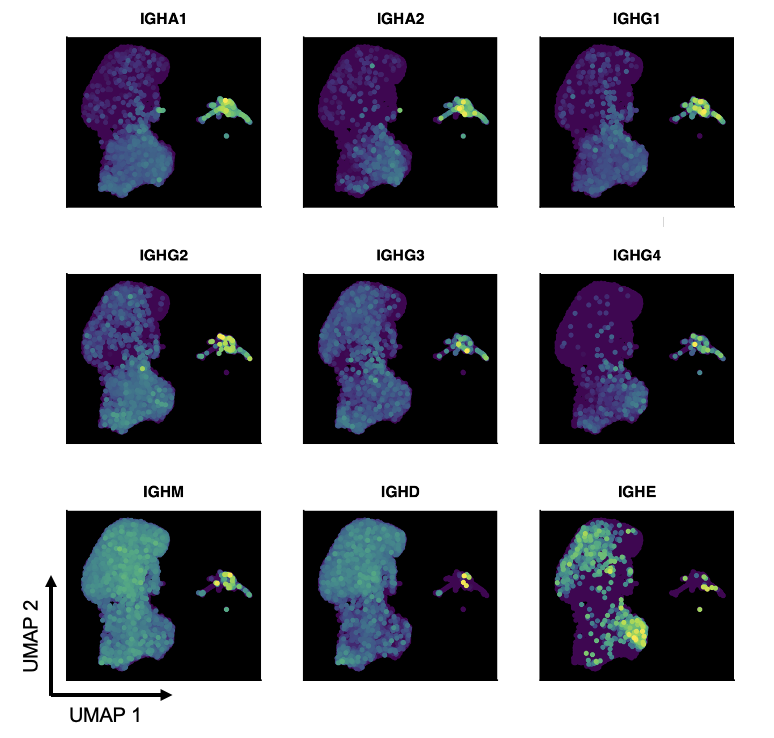

**Figure E9.** B cell feature plots of IgA (*IGHA1, IGHA2*), IgG (*IGHG1, IGHG2, IGHG3, IGHG4*), IgM (*IGHM*), IgD (*IGHD)*, and IgE (*IGHE)*.

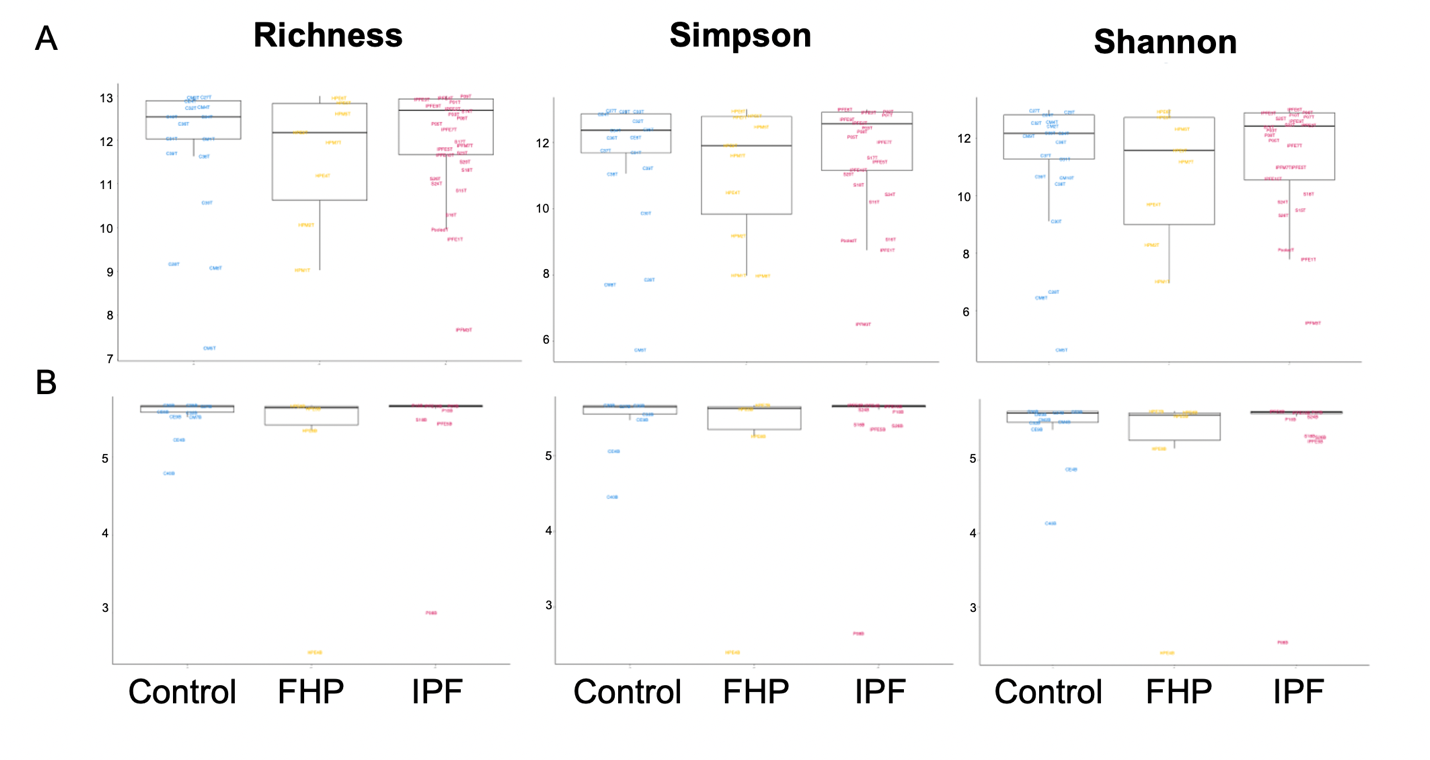

**Figure E10.** Richness, Shannon index, and inverse Simpson diversity indices for the (A) T cell receptor repertoires and (B) B cell receptor repertoires of controls, FHP, and IPF. FHP: fibrotic hypersensitivity pneumonitis; IPF: idiopathic pulmonary fibrosis.

|  | Control (31) | Non-FHP (5) | FHP (14) | IPF (46) |
| --- | --- | --- | --- | --- |
| Age  (mean ± SD) | 68.87 ± 6.06 | 47.00 ± 13.55 | 66.36 ± 11.90 | 70.17 ± 8.01 |
| Sex |  |  |  |  |
| *Male* | 21 | 0 | 6 | 35 |
| *Female* | 10 | 5 | 8 | 11 |
| Ethnicity |  |  |  |  |
| *Caucasian* | 21 | 0 | 6 | 10 |
| *Hispanic* | 10 | 5 | 8 | 34 |
| *Other* | 0 | 0 | 0 | 1 |
| SMOKING |  |  |  |  |
| *NEVER* | 7 | 2 | 8 | 14 |
| *EVER* | 17 | 3 | 6 | 32 |
| *NO DATA* | 7 | 0 | 0 | 0 |
| % FVC (mean ± SD) | 92.00 ± 16.64 | 75.00 ± 15.63 | 61.36 ± 16.95 | 71.88 ± 10.73 |
| % DLCO (mean ± SD) | 104.90 ± 12.80 | 61.00 ± 16.17 | 42.45 ± 15.47 | 50.21 ± 15.23 |

**Table E1. Summary of demographic and clinical characteristics for all processed samples**. Summarized information for patients whose single-cell RNA-sequencing PBMC data were processed for this study. This summary includes all samples before quality control and filtering. Abbreviations: IPF: idiopathic pulmonary fibrosis; FHP: fibrotic hypersensitivity pneumonitis. FVC: Forced Vital Capacity; DLCO: Diffusing Capacity of Carbon Monoxide; SD: Standard Deviation
